# Updated science-wide author databases of standardized citation indicators including retraction data

**DOI:** 10.1101/2024.09.16.613258

**Authors:** John P. A. Ioannidis, Angelo Maria Pezzullo, Antonio Cristiano, Stefania Boccia, Jeroen Baas

**Affiliations:** Department of Medicine, Stanford University, Stanford, California, United States of America; Department of Epidemiology and Population Health, Stanford University, Stanford, California, United States of America; Department of Biomedical Data Science, Stanford University, Stanford, California, United States of America; Department of Statistics, Stanford University, Stanford, California, United States of America; Meta-Research Innovation Center at Stanford (METRICS), Stanford University, Stanford, California, United States of America; Section of Hygiene, Department of Life Sciences and Public Health, Università Cattolica del Sacro Cuore, Rome, Italy; Research Intelligence, Elsevier B.V., Amsterdam, the Netherlands

**Keywords:** retractions, citations, bibliometrics, research practices, meta-research

## Abstract

Citation metrics are widely used in research appraisal, but they provide incomplete views of scientists’ impact and research track record. Other indicators of research practices should be linked to citation data. We have updated a Scopus-based database of highly-cited scientists (top-2% in each scientific subfield according to a composite citation indicator) to incorporate retraction data. Using data from the Retraction Watch database (RWDB), retraction records were linked to Scopus citation data. Of 55,237 items in RWDB as of August 15, 2024, we excluded non-retractions, retractions clearly not due to any author error, retractions where the paper had been republished, and items not linkable to Scopus records. Eventually 39,468 eligible retractions were linked to Scopus. Among 217,097 top-cited scientists in career-long impact and 223,152 in single recent year (2023) impact, 7,083 (3.3%) and 8,747 (4.0%), respectively, had at least one retraction. Scientists with retracted publications had younger publication age, higher self-citation rates, and larger publication volume than those without any retracted publications. Retractions were more common in the life sciences and rare or nonexistent in several other disciplines. In several developing countries, very high proportions of top-cited scientists had retractions (highest in Senegal (66.7%), Ecuador (28.6%) and Pakistan (27.8%) in career-long citation impact lists). Variability in retraction rates across fields and countries suggests differences in research practices, scrutiny, and ease of retraction. Addition of retraction data enhances the granularity of top-cited scientists’ profiles, aiding in responsible research evaluation. However, caution is needed when interpreting retractions, as they do not always signify misconduct; further analysis on a case-by-case basis is essential. The database should hopefully provide a resource for meta-research and deeper insights into scientific practices.

## INTRODUCTION

Citation metrics need to be used with caution (1) to avoid obtaining over-simplified and even grossly misleading views of scientific excellence and impact. The databases of standardized citation metrics across all scientists and scientific disciplines that we have generated using Scopus data (2–4) have attracted wide attention. The databases, which are updated annually (5), have been accessed already more than 3 million times. While providing a centralized, standardized resource may offer an advantage over fragmented, non-standardized efforts, citation data alone offer only a partial view of the impact and work of any given scientist. There is a need to include additional information also on other aspects of their work, including, if possible, further indicators of research quality and integrity (6,7). An important piece of supplemental information to appraise any work is whether a scientist has had publications that have been retracted. In our latest annual update of these citation databases, we have accordingly added information for each included scientist regarding retracted publications.

Retractions are becoming increasingly common, although they still account for a very small proportion of the published literature (8–10). In empirical surveys, various types of misconduct are typically responsible for the majority of retractions (11), and new types of unethical behavior, such as fake papers from paper mills, are becoming increasingly common (12). However, the reasons for retractions are not fully standardized, and many retractions are unclear about why a paper had to be withdrawn. Moreover, some retractions are clearly not due to ethical violations or author errors (e.g. they are due to publisher errors). Finally, in many cases, one may view a retraction as a sign of a responsible author who should be congratulated, rather than chastised, for taking proactive steps to correct the literature. Prompt correction of honest errors, major or minor, is a sign of responsible research practices.

A widely visible list of highly-cited scientists issued annually by Clarivate based on Web of Science no longer includes any scientists with retracted publications (13). In our databases, which cover a much larger number of scientists with more detailed data on each, we have decided to add information on the number of retracted publications, if any, for all listed scientists. Given the variability of the reasons behind retraction, this information can then be interpreted by any assessors on a case-by-case basis with in-depth assessment of reasons and circumstances of each retraction.

Here we present the linkage of retraction and citation data in our latest update of the database of top-cited scientists along with some descriptive statistics related to this new combined resource. We hope that this new resource will be useful for further meta-research studies that may be conducted by investigators on diverse samples of scientists and scientific fields.

## METHODS AND RESULTS

To add the new information on retractions, we depended on the most reliable database of retractions available to date, the Retraction Watch database (RWDB) (13) which is also publicly freely available through CrossRef. Among the 55,237 RWDB entries obtained from CrossRef (https://api.labs.crossref.org/data/retractionwatch) on August 15, 2024, we focused on the 50,457 entries where the nature of the notice is classified as "Retraction", excluding other types (corrections, expressions of concern) that may also be covered in RWDB. From this set, we excluded entries where the paper had been retracted but then replaced by a new version (which can suggest that the errors were manageable to address and there is a new version representing the work in the published literature); and those entries where the retraction was clearly solely not due to any error or wrong-doing by the authors (e.g. publisher error). Therefore, we excluded entries where the reason for retraction was listed as "Retract and Replace," "Error by Journal/Publisher," "Duplicate Publication through Error by Journal/Publisher", or "Withdrawn (out of date)"; however, for the latter three categories, these exclusions were only applied if there were no additional reasons listed that could be attributed potentially to the authors exclusively or in part, as detailed in Supplementary Table 1. This resulted in a set of 47,964 entries.

Following this filtering process, we linked the retraction records to Scopus using the digital object identifier (DOI) of the original paper and this resulted in 38,364 matches. For entries where a DOI match was not possible, we attempted to link records using a combination of the title and year derived from the date of the original article, allowing for a +/- 1-year variation, resulting in 1,104 additional matches. This process resulted in a total of 39,468 matched records.

Calculation of the composite citation indicator and ranking of the scientists accordingly within their primary subfield (using the Science-Metrix classification of 20 fields and 174 subfields) were performed in the current iteration with the exact same methods as in previous iterations (described in detail in references (2–4)).

The new updated release of the databases includes 217,097 scientists who are among the top-2% of their primary scientific subfield in the career-long citation impact and 223,152 scientists who are among the top-2% in their single most recent year (2023) citation impact. These numbers also include some scientists (2,789 and 6,325 scientists in the two datasets, respectively) who may not be in the top-2% of their primary scientific subfield but are among the 100,000 top-cited across all scientific subfields combined. Among the top-cited scientists, 7,083 (3.3%) and 8,747 (4.0%), respectively, in the two datasets have at least one retracted publication. 1,710 (0.8%) and 2,150 (1.0%), respectively, have 2 or more retracted publications. As shown in Figure 1, the distribution of the number of linked eligible retractions per author follows a power law.

**Figure 1.**
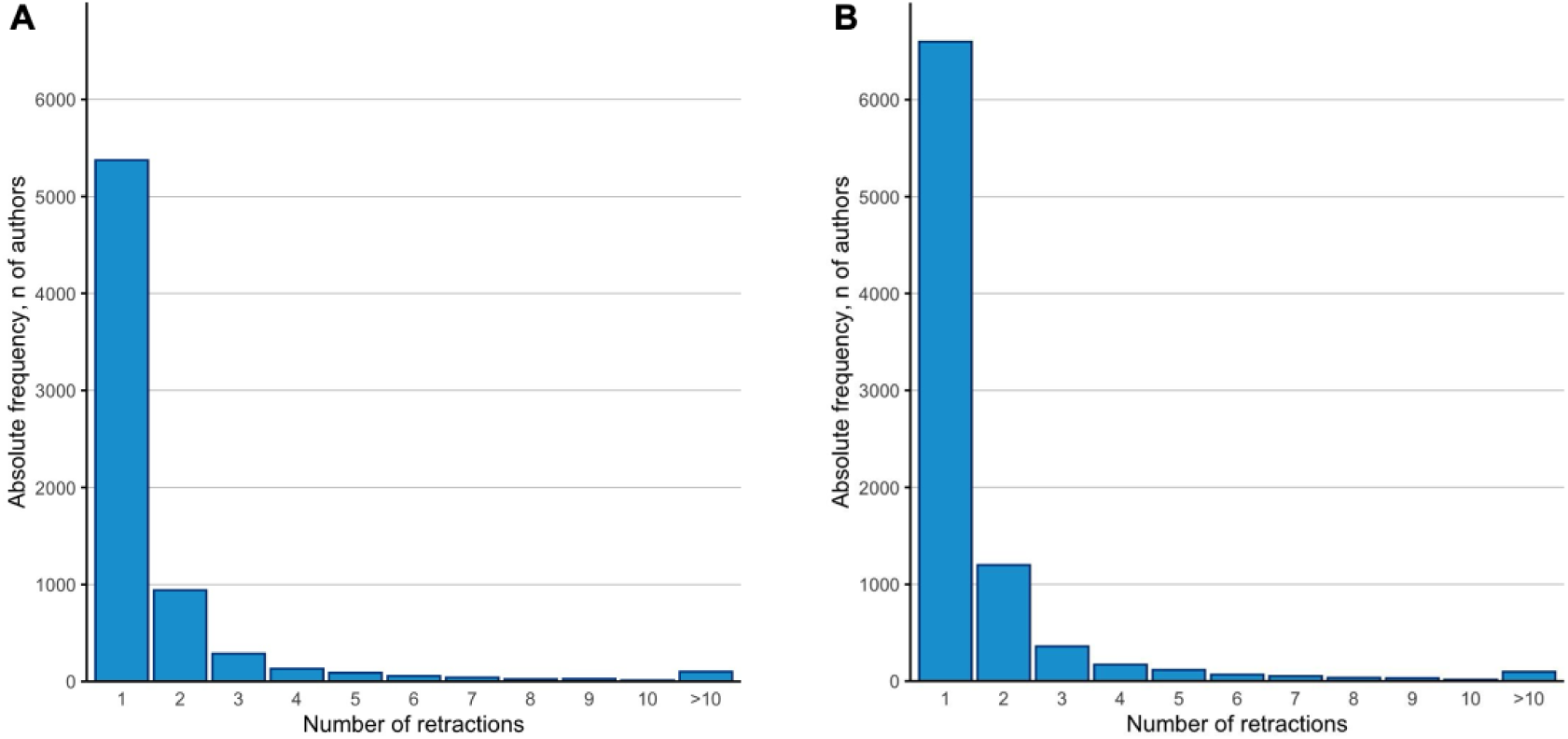
Distribution of the number of retractions in top-cited scientists with at least one retraction. A Database of top-cited authors based on career-long impact. B. Database of top-cited authors based on single recent year (2023) impact.

Table 1 shows the characteristics of those top-cited scientists who have any retracted publications versus those who have not had any retractions. As shown, top-cited scientists with retracted publications tend to have younger publication ages, higher proportion of self-citations, higher ratio of h/hm index (indicating higher co-authorship levels), slightly better ranking, and higher total number of publications (p<0.001 by Mann-Whitney *U* test for all indicators in the career-long impact dataset and the single recent year dataset, except for the publication age and the absolute ranking in the subfield in the single recent year dataset). However, except for the number of papers published, the differences are small or modest in absolute magnitude. The proportion of scientists with retractions is highest though at the extreme top of ranking. Among the top-1000 scientists with the highest composite indicator values, the proportion of those with at least one retraction are 13.8% and 11.1%, in the career-long and single recent year impact, respectively.

**Table 1.**
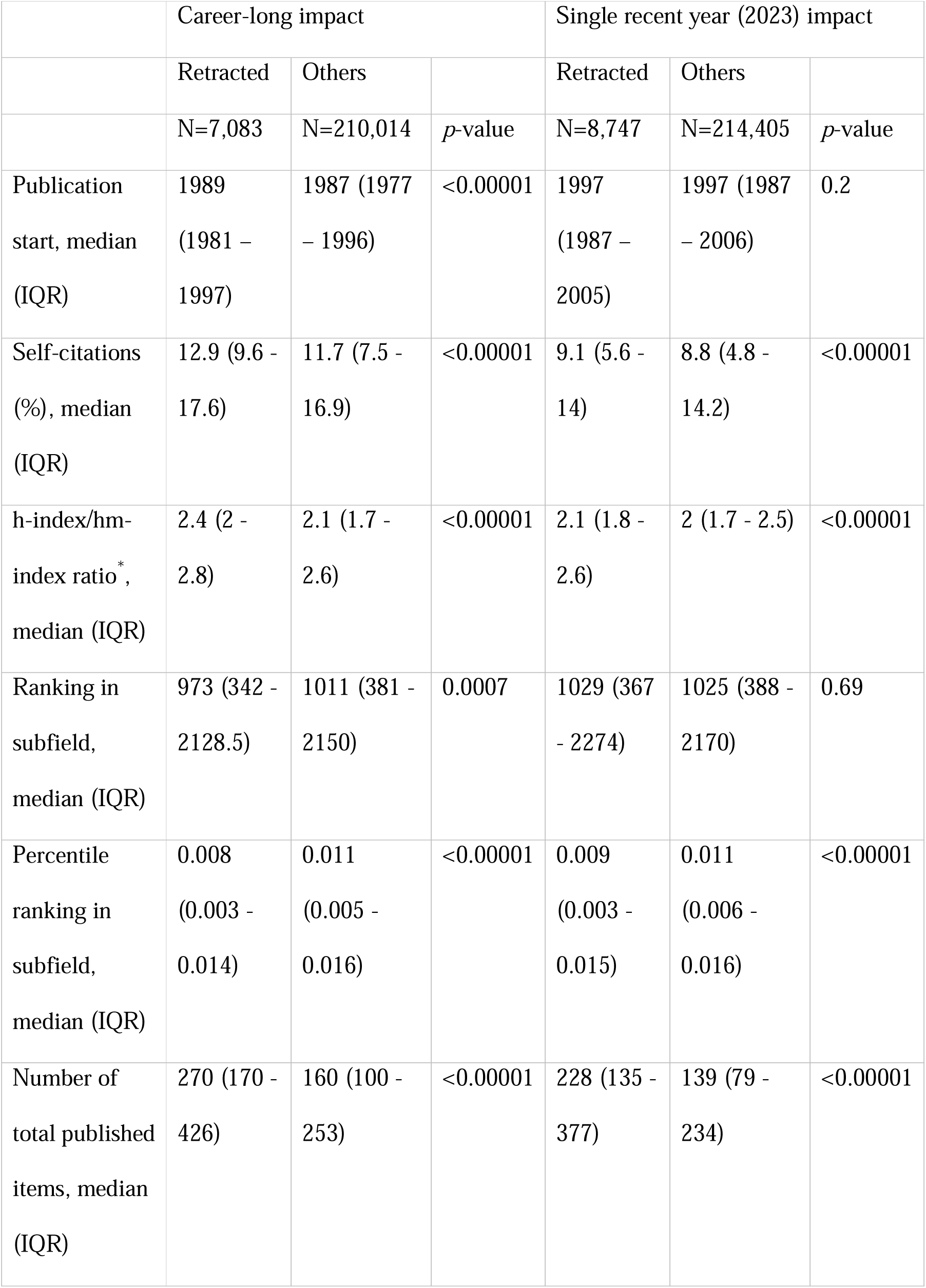
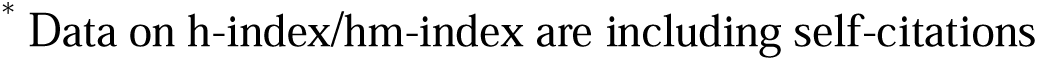
Top-cited scientists with and without retracted publications characteristics and Mann-Whitney *U* test.

|  | Career-long impact |  |  | Single recent year (2023) impact |  |  |
| --- | --- | --- | --- | --- | --- | --- |
|  | Retracted | Others |  | Retracted | Others |  |
|  | N=7,083 | N=210,014 | *p*-value | N=8,747 | N=214,405 | *p*-value |
| Publication start, median (IQR) | 1989 (1981 – 1997) | 1987 (1977 – 1996) | <0.00001 | 1997 (1987 – 2005) | 1997 (1987 – 2006) | 0.2 |
| Self-citations (%), median (IQR) | 12.9 (9.6 - 17.6) | 11.7 (7.5 - 16.9) | <0.00001 | 9.1 (5.6 - 14) | 8.8 (4.8 - 14.2) | <0.00001 |
| h-index/hm-index ratio*, median (IQR) | 2.4 (2 - 2.8) | 2.1 (1.7 - 2.6) | <0.00001 | 2.1 (1.8 - 2.6) | 2 (1.7 - 2.5) | <0.00001 |
| Ranking in subfield, median (IQR) | 973 (342 - 2128.5) | 1011 (381 - 2150) | 0.0007 | 1029 (367 - 2274) | 1025 (388 - 2170) | 0.69 |
| Percentile ranking in subfield, median (IQR) | 0.008 (0.003 - 0.014) | 0.011 (0.005 - 0.016) | <0.00001 | 0.009 (0.003 - 0.015) | 0.011 (0.006 - 0.016) | <0.00001 |
| Number of total published items, median (IQR) | 270 (170 - 426) | 160 (100 - 253) | <0.00001 | 228 (135 - 377) | 139 (79 - 234) | <0.00001 |

\* Data on h-index/hm-index are including self-citations

Table 2 shows the proportion of top-cited scientists with retracted publications across the 20 major fields that science is divided according to the Science-Metrix classification; information on the more detailed 174 subfields appears in Supplementary Table 2. The proportion of retractions varies widely across major fields, ranging from 0% to 5.6%. Clinical Medicine and Biomedical Research have the highest rates (4.9-5.6%). Enabling & Strategic Technologies, Chemistry and Biology have rates close to the average of all sciences combined. All other fields have from low to very low (or even zero) rates of scientists with retractions. When the 174 Science-Metrix subfields of science were considered, the highest proportions of top-cited scientists with at least one retracted paper were seen in the subfields of Complementary & Alternative Medicine, Oncology & Carcinogenesis and Pharmacology & Pharmacy (with 10.5%, 9.9%, and 9.4%, respectively of top-cited scientists based on single recent year impact). See details on all 174 subfields in Supplementary Table 2.

**Table 2.**
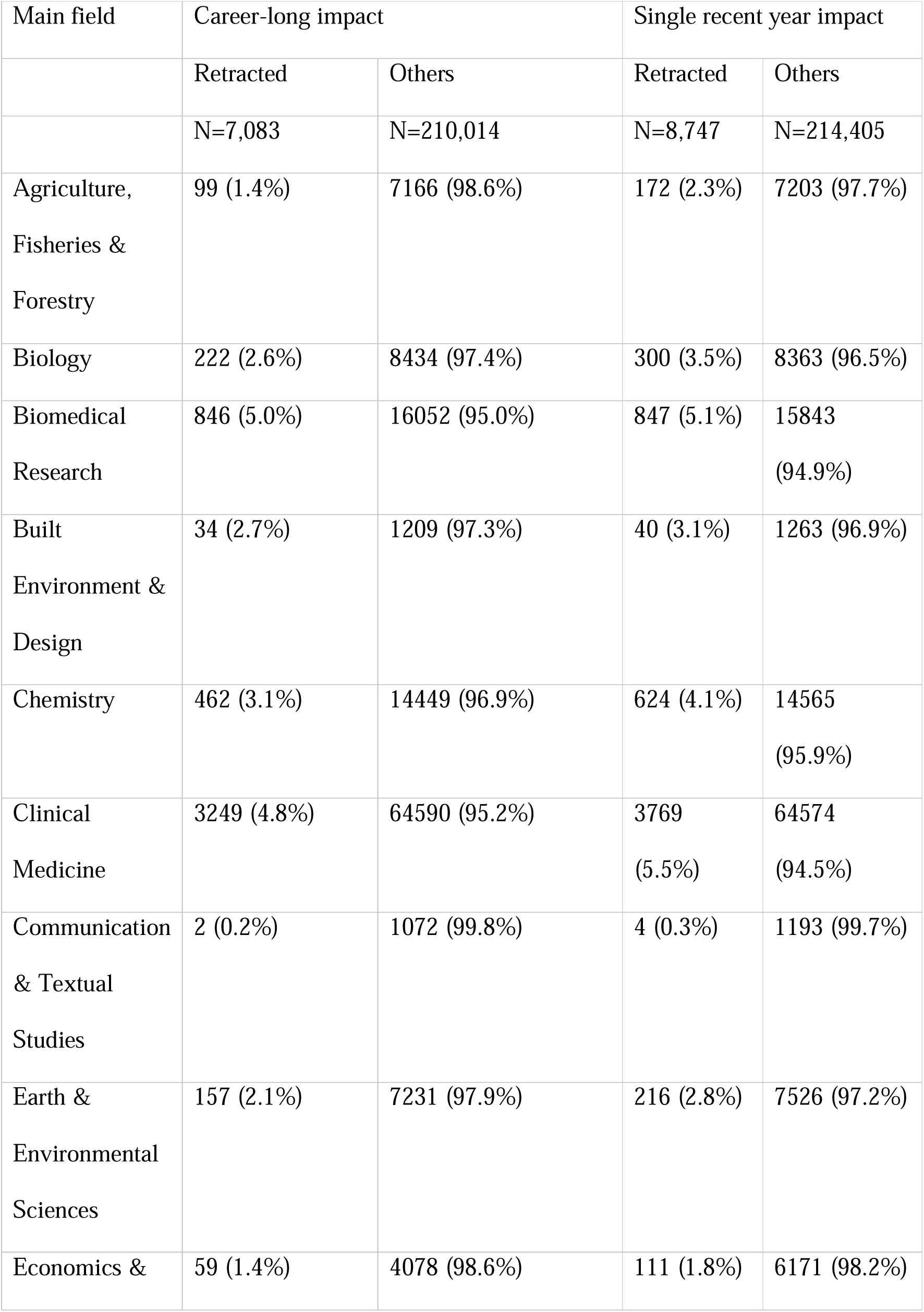
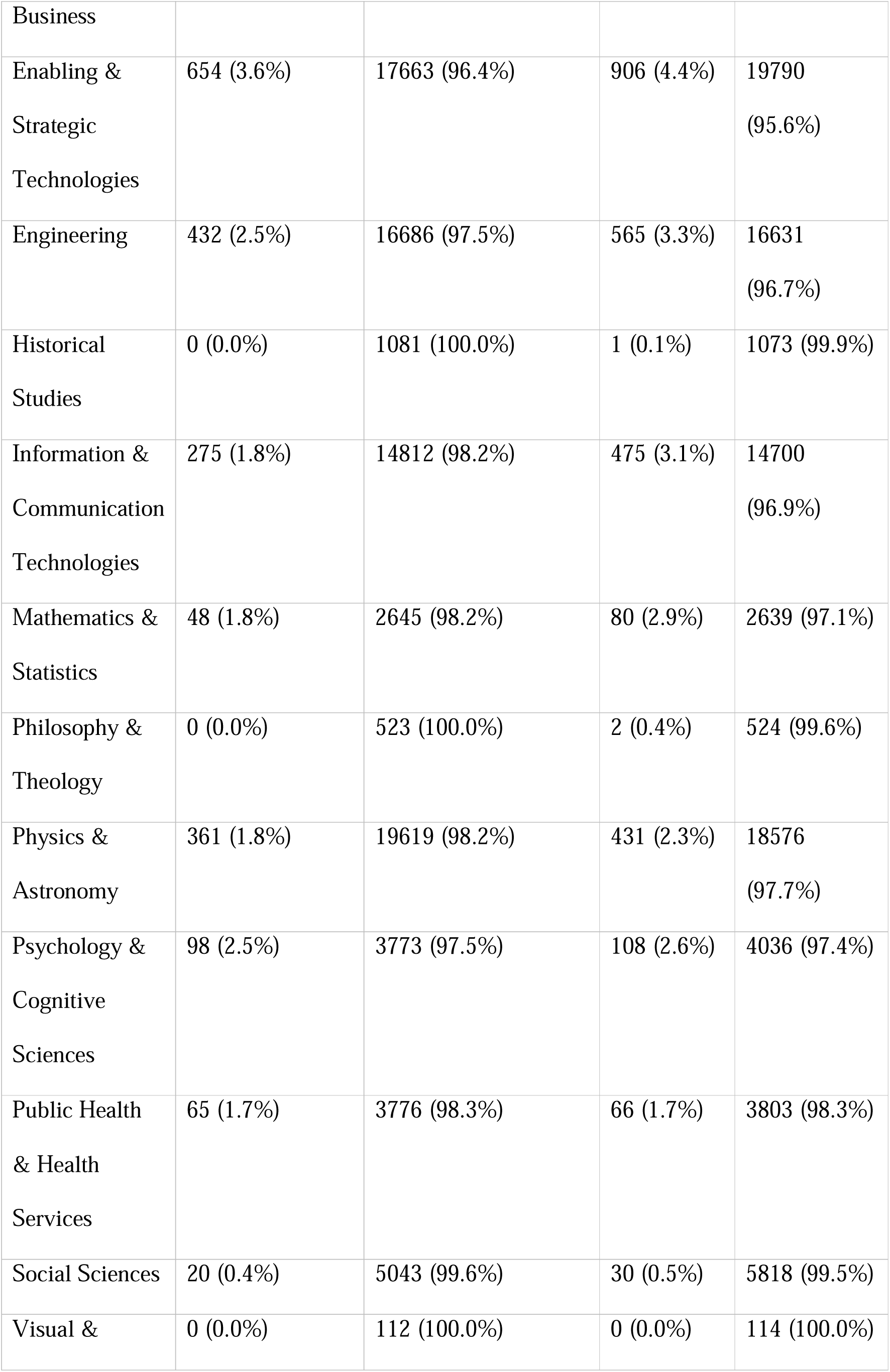
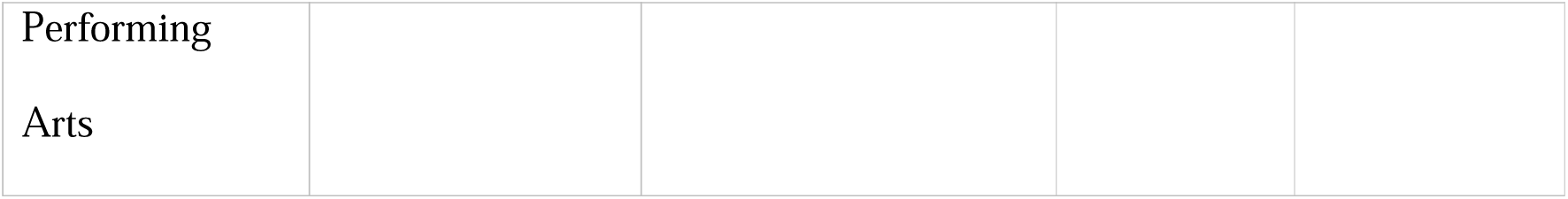
Top-cited scientists with and without/ retracted publications according to their main field.

Retraction rates among top-cited scientists also vary in the 20 countries that host most of the top-cited authors (Table 3), with higher rates observed in India (9.2% career-long - 8.6% single recent year impact), China (8.2% - 6.7%) and Taiwan (5.2% - 5.7%), and lower rates observed in Israel (1.7% – 2.0%), Belgium (2.1% – 2.1%), and Finland (2.2% - 2.2%). Some countries with few top-cited authors (not among the 20 shown in Table 2) have impressive rates of scientists with retractions: Countries that exceed 10% either in career-long or in single recent year top-cited scientists are listed in Supplementary Table 3. The highest proportions of top-cited scientists with retractions were seen in Senegal (66.7%), Ecuador (28.6%), and Pakistan (27.8%) based on the career-long impact list and in Kyrgyzstan (50%), Senegal (41.7%), Ecuador (28%) and Belarus (26.7%) in the single recent year impact list.

**Table 3.** Top-cited scientists with and without retracted publications according to country.

| Country | Career-long impact |  | Single recent year impact |  |
| --- | --- | --- | --- | --- |
|  | Retracted | Others | Retracted | Others |
| United States | 2332 (2.8%) | 81870 (97.2%) | 2186 (3.1%) | 69206 (96.9%) |
| United Kingdom | 430 (2.2%) | 19218 (97.8%) | 428 (2.4%) | 17127 (97.6%) |
| Germany | 336 (2.9%) | 11236 (97.1%) | 309 (3.0%) | 10111 (97.0%) |
| China | 877 (8.2%) | 9810 (91.8%) | 1813 (6.7%) | 25352 (93.3%) |
| Canada | 241 (2.6%) | 9024 (97.4%) | 223 (2.7%) | 7962 (97.3%) |
| Japan | 362 (4.4%) | 7899 (95.6%) | 254 (4.5%) | 5354 (95.5%) |
| Australia | 178 (2.4%) | 7270 (97.6%) | 201 (2.5%) | 7833 (97.5%) |
| France | 151 (2.2%) | 6770 (97.8%) | 152 (2.6%) | 5630 (97.4%) |
| Italy | 254 (4.1%) | 6017 (95.9%) | 300 (3.9%) | 7318 (96.1%) |
| Netherlands | 123 (2.7%) | 4392 (97.3%) | 116 (2.6%) | 4419 (97.4%) |
| Spain | 103 (2.9%) | 3405 (97.1%) | 127 (3.2%) | 3880 (96.8%) |
| Switzerland | 84 (2.4%) | 3347 (97.6%) | 82 (2.4%) | 3323 (97.6%) |
| Sweden | 84 (2.5%) | 3269 (97.5%) | 78 (3.0%) | 2566 (97.0%) |
| India | 270 (9.2%) | 2669 (90.8%) | 462 (8.6%) | 4889 (91.4%) |
| South Korea | 120 (5.1%) | 2246 (94.9%) | 186 (5.3%) | 3313 (94.7%) |
| Denmark | 46 (2.2%) | 2068 (97.8%) | 53 (2.6%) | 1960 (97.4%) |
| Israel | 36 (1.7%) | 2057 (98.3%) | 32 (2.0%) | 1590 (98.0%) |
| Belgium | 41 (2.1%) | 1956 (97.9%) | 42 (2.1%) | 1965 (97.9%) |
| Taiwan | 91 (5.2%) | 1668 (94.8%) | 80 (5.7%) | 1327 (94.3%) |
| Finland | 32 (2.2%) | 1413 (97.8%) | 26 (2.2%) | 1153 (97.8%) |

The new iteration of the two top-cited scientists’ datasets also includes information on the number of citations received (overall and in the single recent year, respectively) by the retracted papers of each scientist. If we consider scientists with at least one retraction, the range is 0 to 7,491, with median (IQR) of 25 (6 to 80) in the career-long dataset. The range is 0 to 832 with median (IQR) of 1 (0 to 4) in the single recent year dataset. A total of 114 scientists in the career-long dataset have received more than 1,000 citations to their retracted papers and for 230 (0.1%) and 260 (0.1%) scientists in the two datasets the citations to the retracted papers account for more than 5% of their citations.

Furthermore, information is provided for each scientist on the number of citations that they received from any of the 39,950 retracted papers. In the career-long dataset, the range is 0 to 1,974, with median (IQR) of 0 (2 to 5) and in the single recent year dataset, the range is 0 to 180 with median (IQR) of 0 (0 to 0). A total of 5 scientists in the career-long dataset have received more than 1,000 citations from papers that have been retracted and for 14 and 7 scientists in the two datasets, the citations they have received from retracted papers account for more than 5% of their citations (overall and in the single recent year, respectively).

## DISCUSSION

We hope that the addition of the retraction data will improve the granularity of the information provided on each scientist in the new, expanded database of top-cited scientists. A more informative profile may be obtained by examining not only the citation indicators but retracted papers, proportion of self-citations, evidence of extremely prolific behavior (14) (see detailed data that can be linked to the top-cited scientists’ database, published in https://elsevier.digitalcommonsdata.com/datasets/kmyvjk3xmd/2), as well as responsible indicators such as data and code sharing and protocol registration information that is becoming increasingly available (6,7).

The data suggest that approximately 4% of the top-cited scientists have at least one retraction. This is a conservative estimate, and the true rate may be higher since some retractions are in titles that are not covered by Scopus or could not be linked in our dataset linkage. Proportions of scientists with retractions are substantially higher in the extremes of the most-cited scientists. Top-cited scientists with retracted publications exhibit higher levels of collaborative co-authorship and have a higher total number of papers published. High productivity and more extensive co-authorship may be associated with less control over what gets published or may show proficiency in gaming the system. Nevertheless, the higher publication output of scientists with retractions might simply reflect that the more you publish, the greater the chance of encountering eventually a retraction.

Retractions are far more common in the life sciences than in other fields. Many scientific fields have minimal or no track records of retractions and some subfields such as alternative medicine, cancer research, and pharmacology exhibit rates of retractions double the rates exhibited by the life sciences overall. These differences might reflect the increased scrutiny and better detection of misconduct and major errors in fields that have consequences for health; differences in the intensity and types of post-publication review practices (15); and the fact that quantifiable data and images in the life sciences are easier to assess for errors and fraud than many constructs in social sciences.

Many developing countries have extremely high rates of top-cited authors with retracted papers. This may reflect problematic research environments and incentives in these countries, several of which are also rapidly growing their overall productivity (10,16–19). In fact, some of these countries such as India, China, Pakistan and Iran also have a large share of implausibly hyperprolific authors (14). It would be interesting to see if removing some of the productivity incentives may reduce the magnitude of the problem in these countries.

As previously documented, several retracted papers have been cited considerably and, unfortunately, some continue to be cited even after their retraction (20,21). This is a problem that should and can be hopefully fixed.

Among top-cited authors, a small number have received a very large number of citations to their retracted papers. However, these citations have a relatively small proportional contribution to the overall very high total citation counts of these scientists. The same applies to the proportion of citations that are received by retracted papers. Some highly-cited authors may have received a substantial number of citations from retracted papers, but this is a very small proportion against their total citations. Nevertheless, within paper mills, fake papers may be using repeatedly the same citations from known, influential authors and papers that are already cited heavily in the literature. It is possible that most paper mill products remain undetected and have not yet been retracted from the literature.

We expect that the new, expanded database may enhance the progression of further research on citation and retraction indicators, with expanded linkage to yet more research indicators. We caution that even though we excluded retractions that attributed no fault to the authors, we cannot be confident that all the included retractions included some error, let alone misconduct, by the authors. Moreover, sometimes not all authors may have been responsible for what led to the retraction. Therefore, any further analyses that focus on individual author profiles rather than aggregate, group-level analyses should pay due caution in dissecting the features and circumstances surrounding each retraction. Unfortunately, these are often not presented in sufficient detail to allow safe judgements (11,22).

Moreover, inaccuracies are possible in the merged dataset. As discussed previously, Scopus has high precision and recall (23), but some errors do exist in author ID files. Errors may also happen in the attribution of affiliation for each scientist. Finally, considering the vast size of these datasets with potential duplicity and similarity of names, ensuring that no scientist is incorrectly associated with a retracted paper is virtually impossible. Users of these datasets and/or Scopus can improve author profile accuracy by offering corrections directly to Scopus through the use of the Scopus to ORCID feedback wizard (https://orcid.scopusfeedback.com/). Most importantly, we make no judgment calls in our databases on the ethical nature of the retractions; similarly we do not comment on whether the retractions may be fair or not. Some retractions may still be contested between authors and editors and/or may even have ongoing legal proceedings. We urge users of these data to very carefully examine the evidence and data surrounding each retraction and its nature.

## Acknowledgments

This work uses Scopus data provided by Elsevier. We are thankful to Alison Abritis and Ivan Oransky for constructive comments. The work of Antonio Cristiano in this research has been supported by the European Network Staff Exchange for Integrating Precision Health in the Healthcare Systems project (Marie Skłodowska-Curie Research and Innovation Staff Exchange no. 823995).

---

**Supplementary Table 1.**
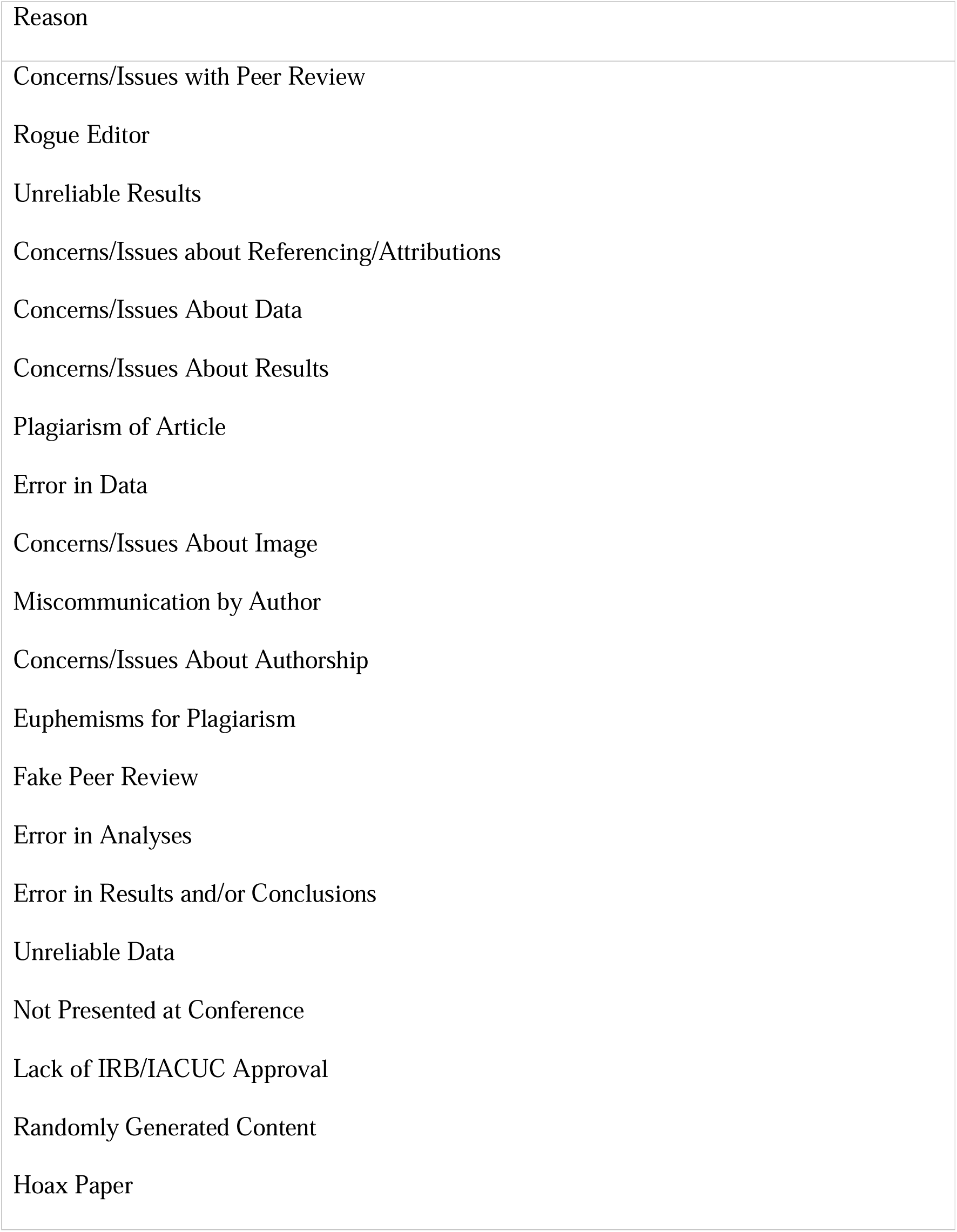
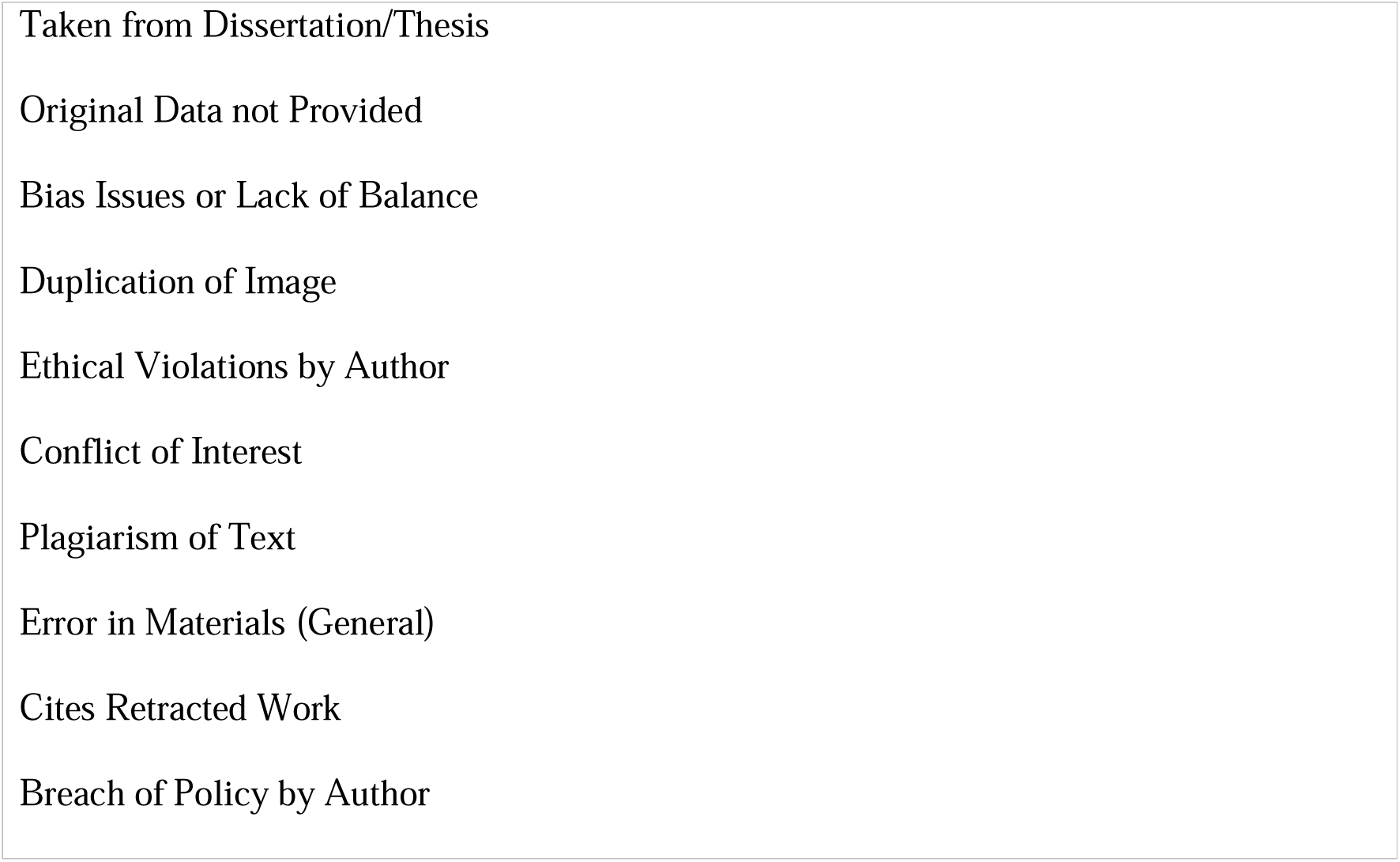
List of author-attributable reasons used to filter journal error and withdrawn (out of date) exceptions. For a full description of these reasons, please refer to: https://retractionwatch.com/retraction-watch-database-user-guide/retraction-watch-database-user-guide-appendix-b-reasons/.

**Supplementary Table 2.**
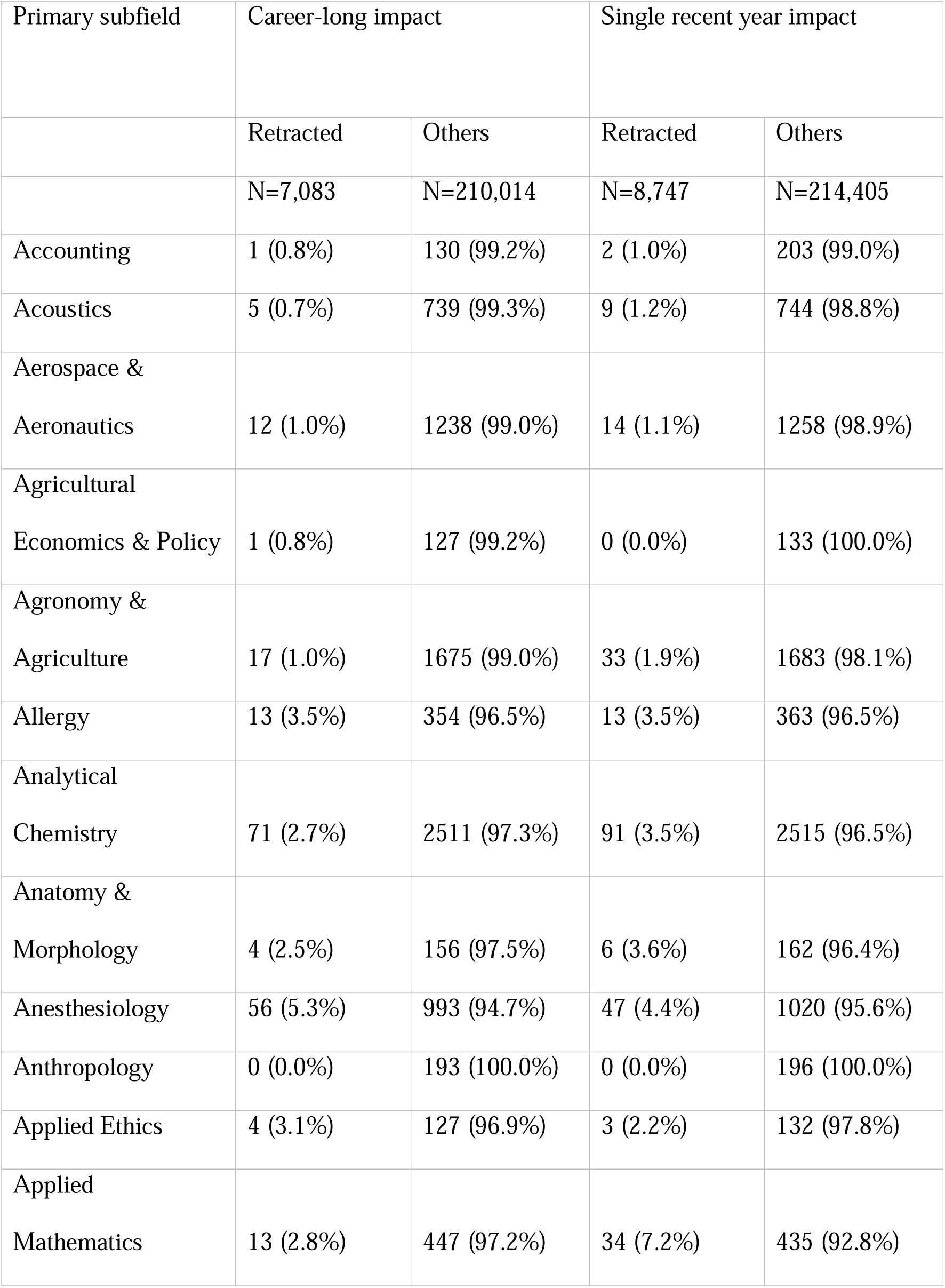
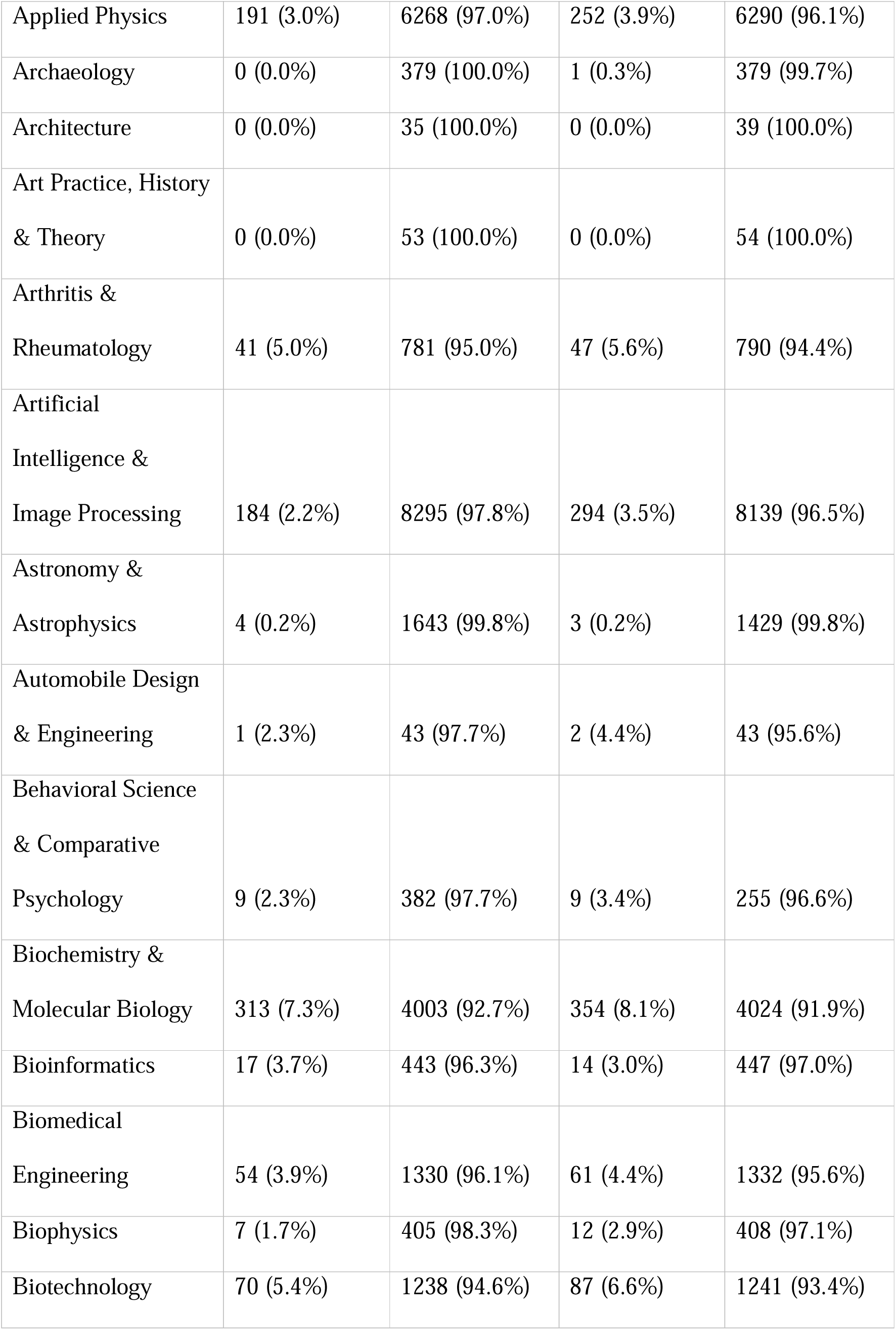
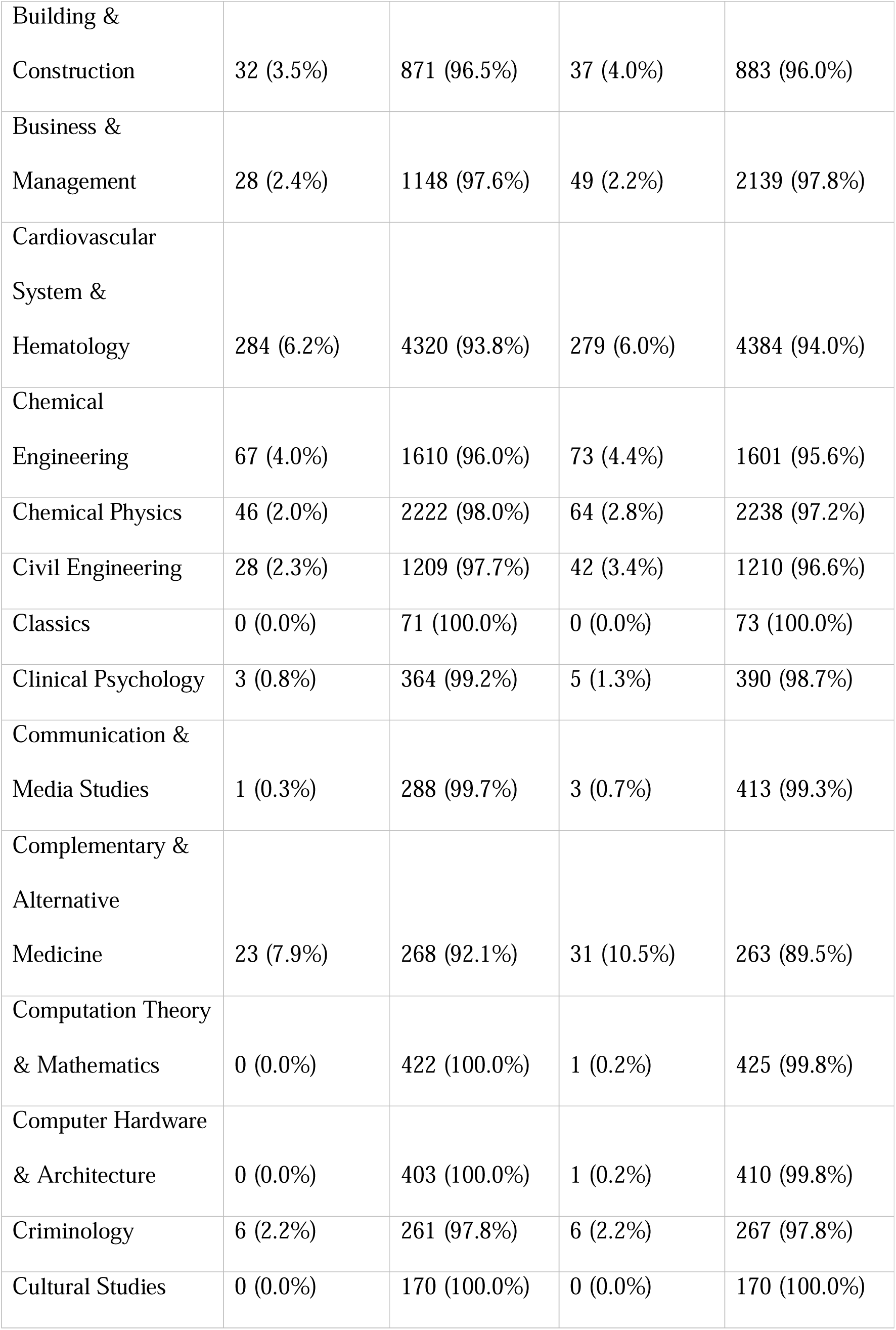
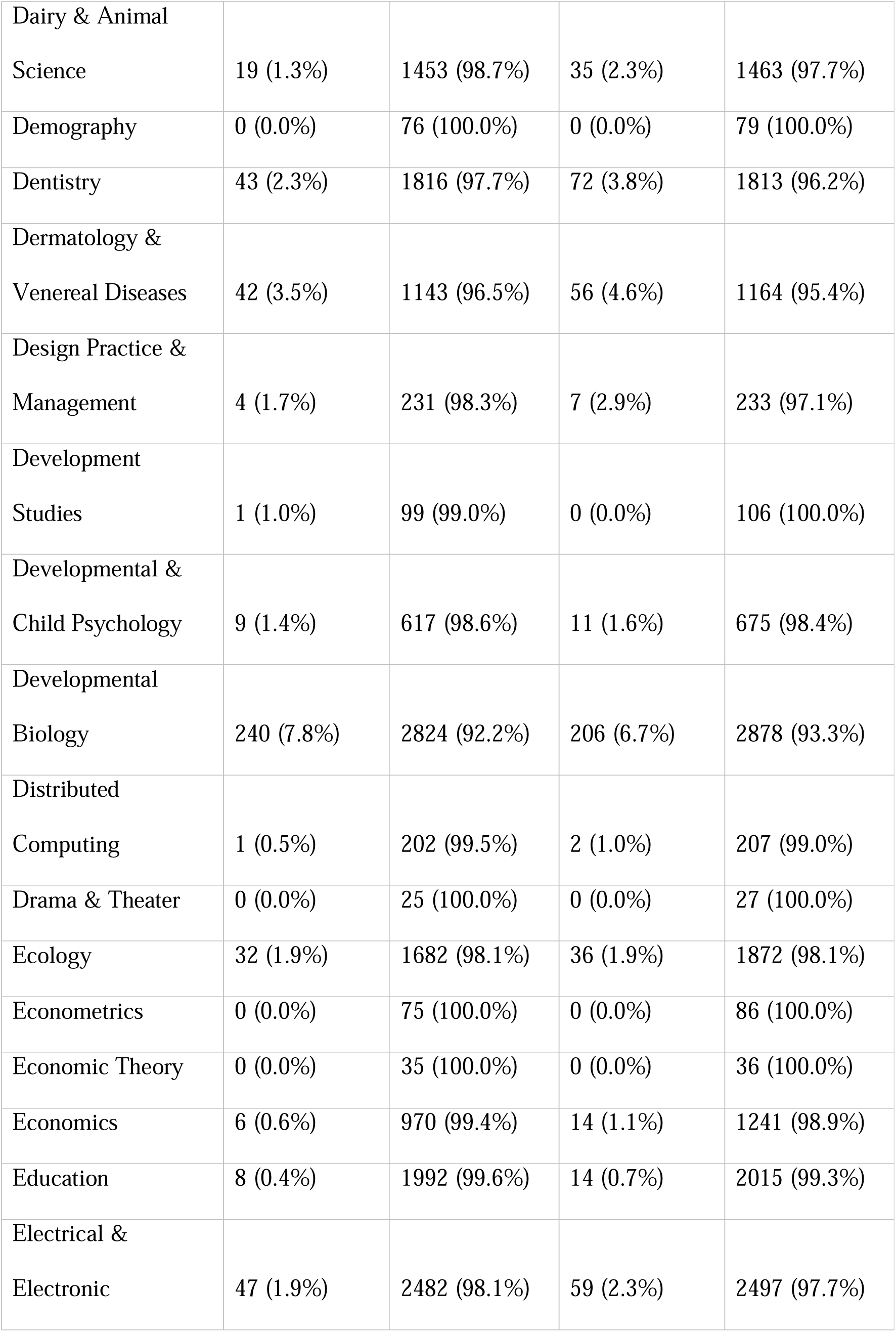
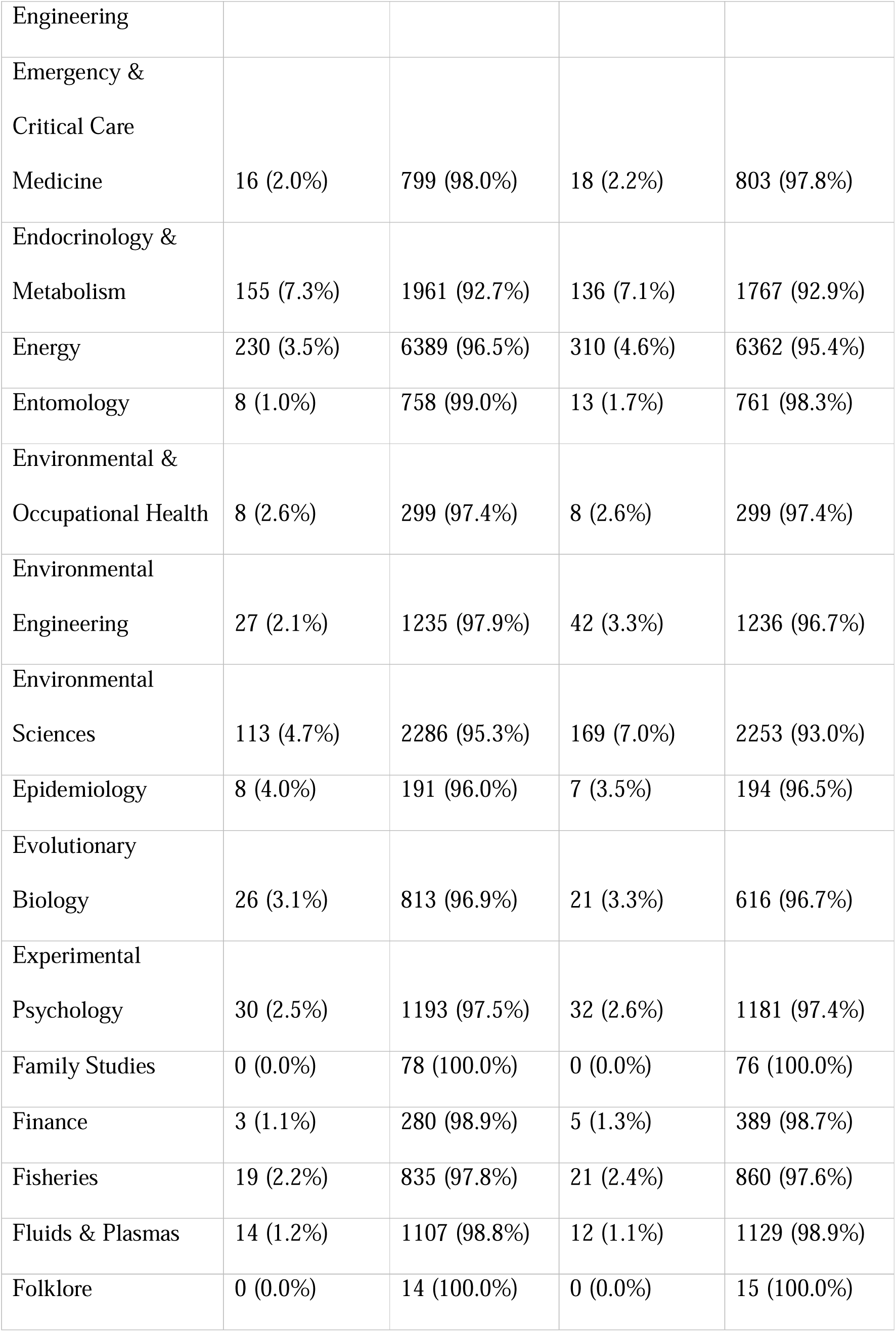
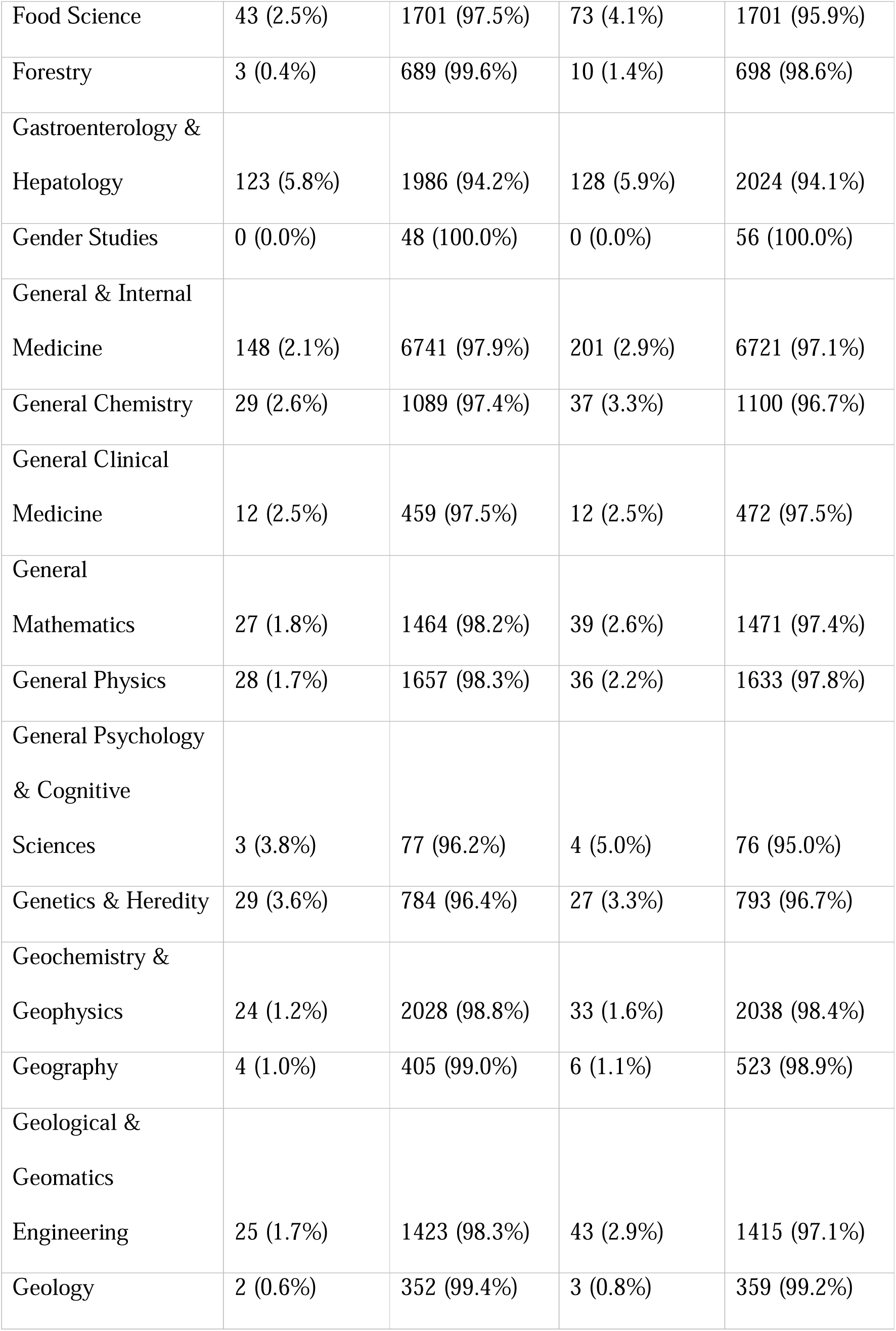
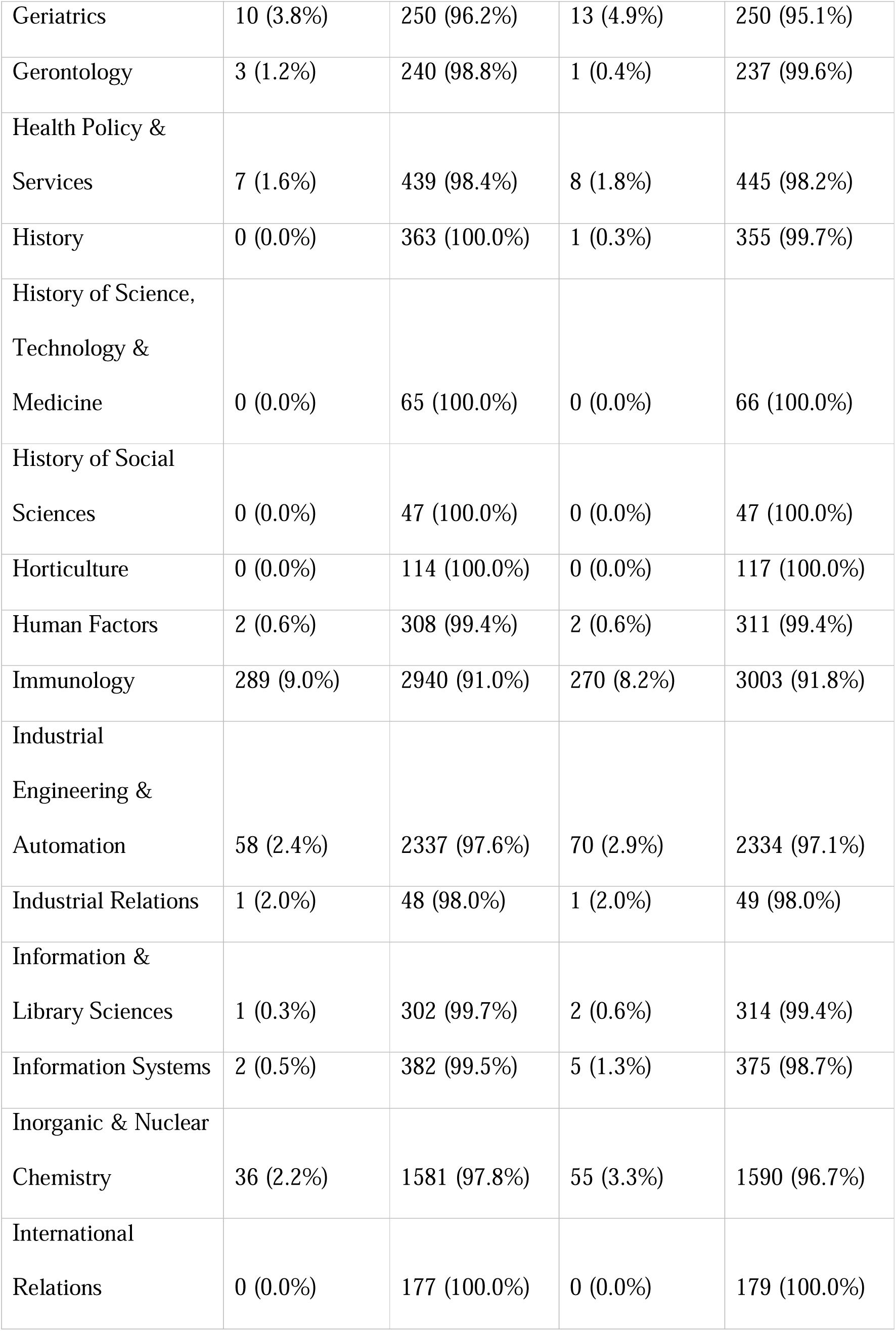
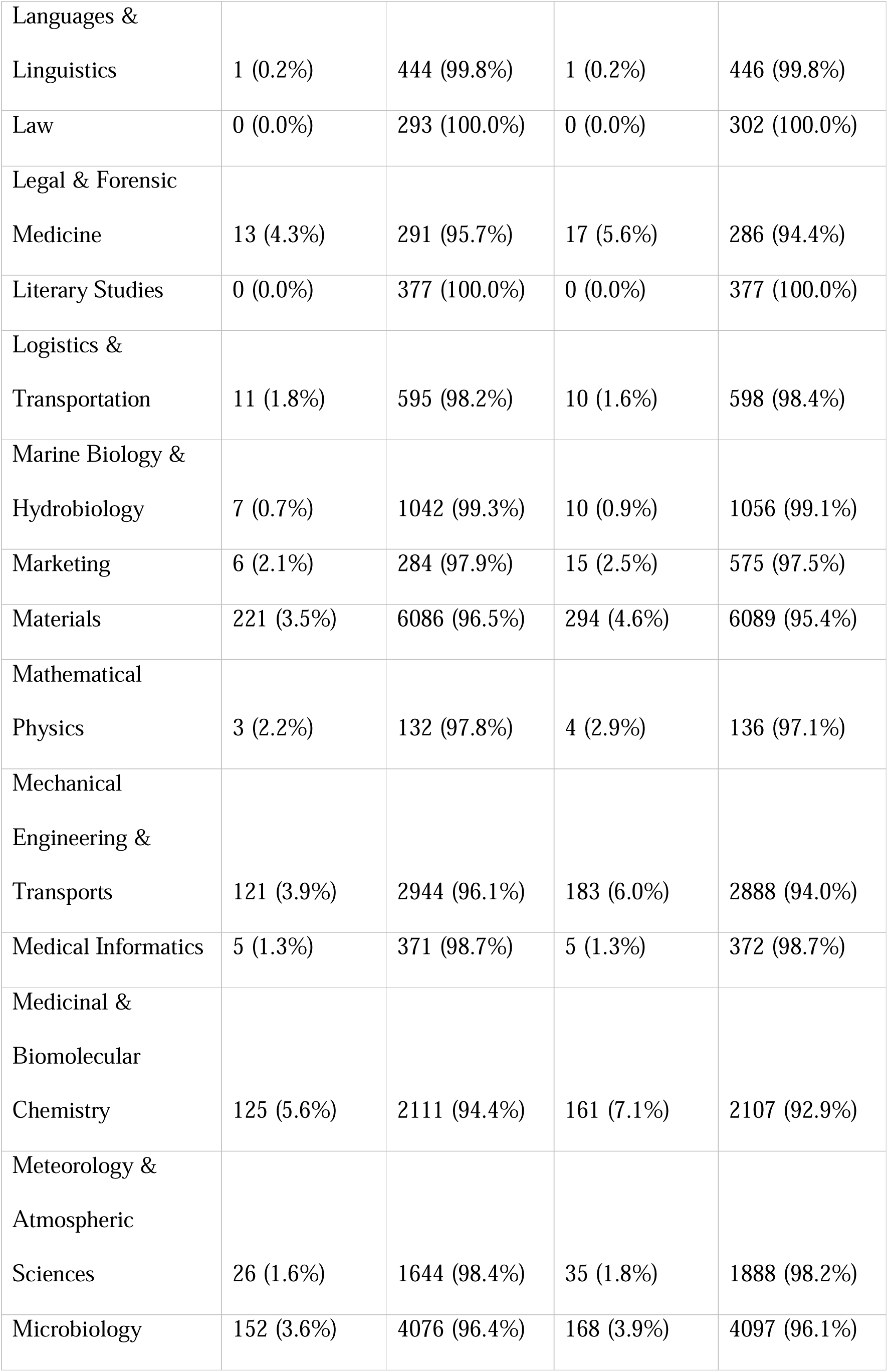
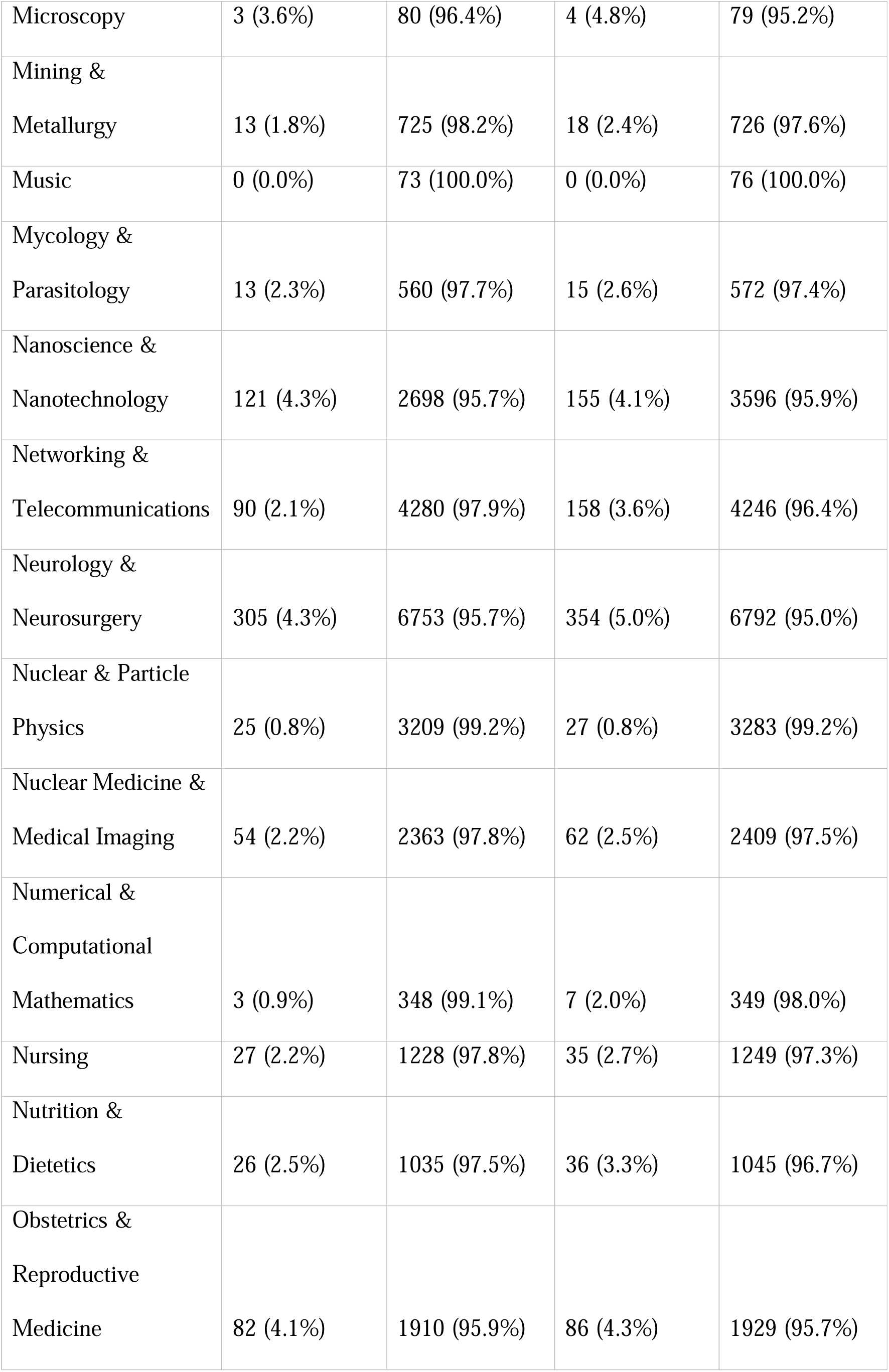
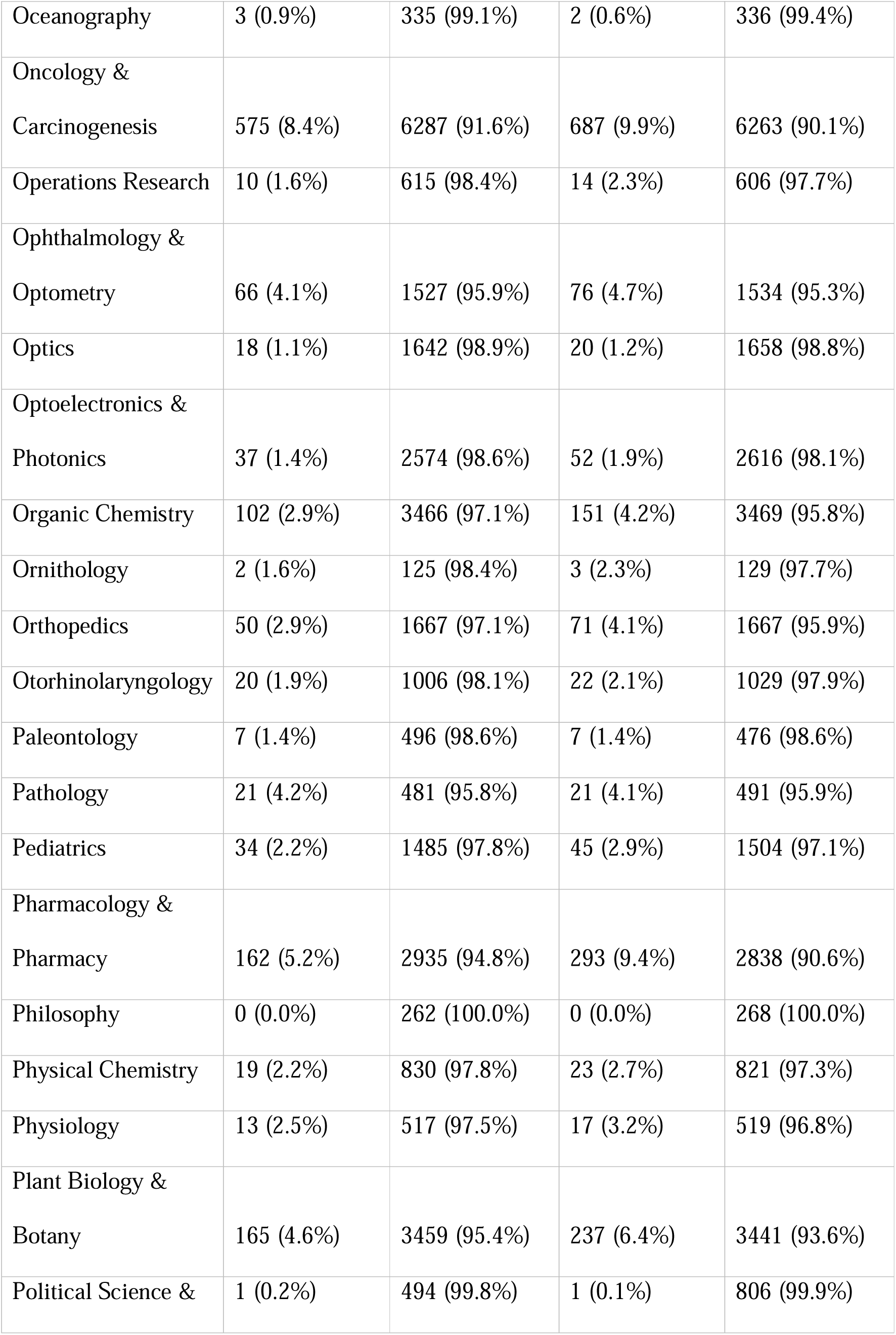
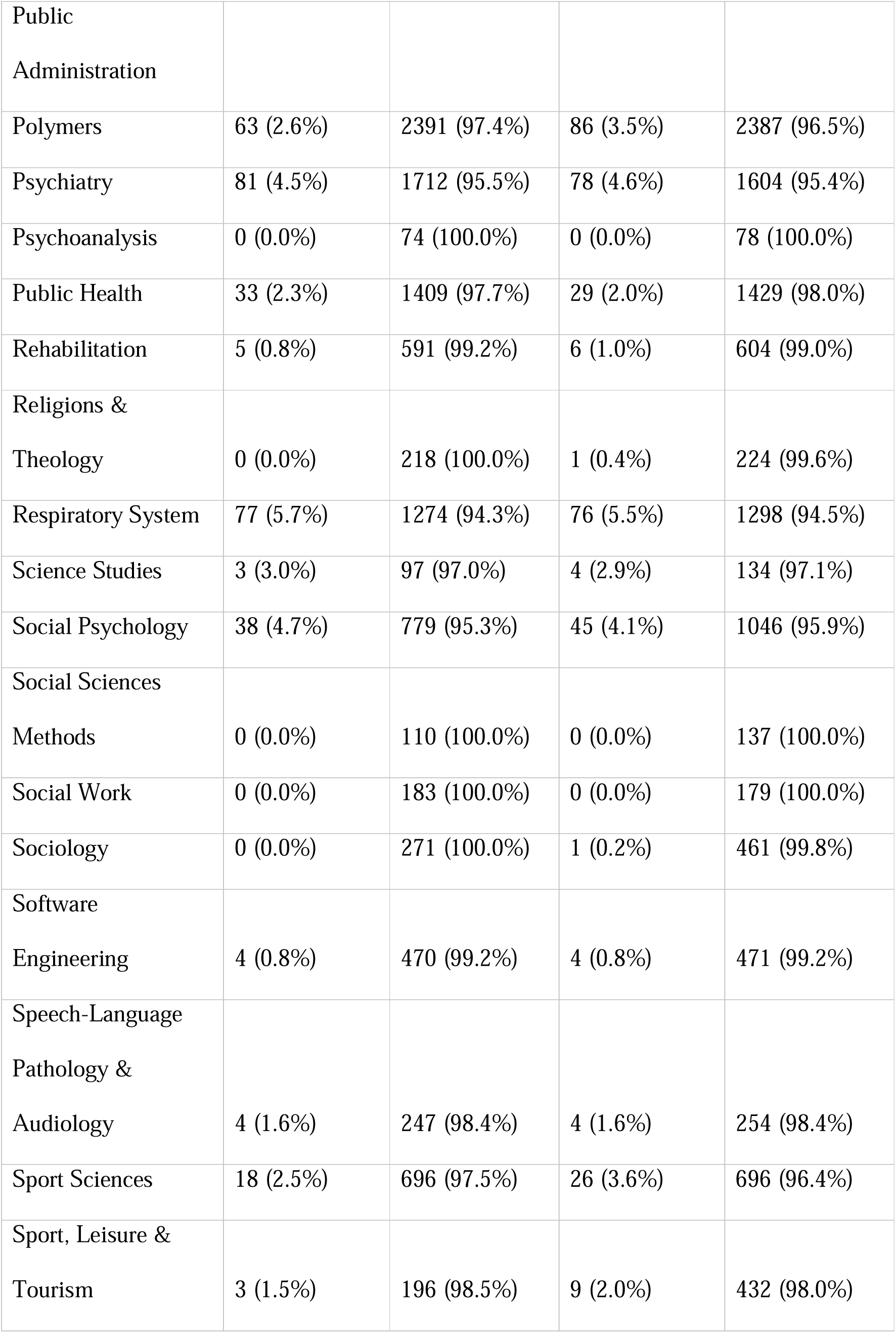
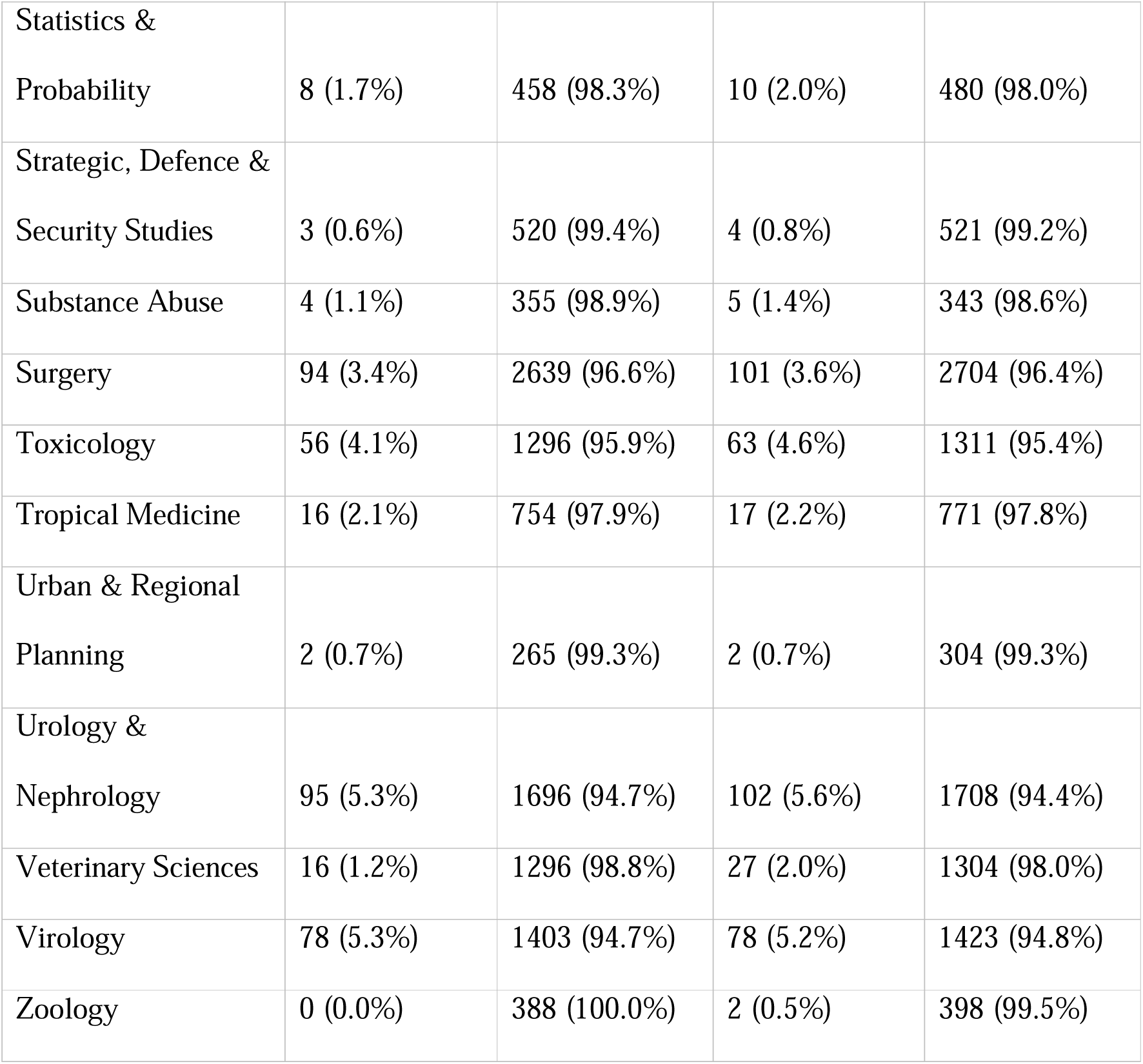
Top-cited scientists with and without retracted publications according to their primary subfield.

**Supplementary Table 3.** Top-cited scientists with and without retracted publications in countries with high (>10%) retraction prevalence.

| Country | Career-long impact |  | Single recent year impact |  |
| --- | --- | --- | --- | --- |
|  | Retracted | Others | Retracted | Others |
| Armenia | 0 (0.0%) | 7 (100.0%) | 1 (11.1%) | 8 (88.9%) |
| Azerbaijan | 1 (11.1%) | 8 (88.9%) | 2 (11.1%) | 16 (88.9%) |
| Bangladesh | 8 (11.8%) | 60 (88.2%) | 19 (9.3%) | 186 (90.7%) |
| Belarus | 2 (8.7%) | 21 (91.3%) | 4 (26.7%) | 11 (73.3%) |
| Ecuador | 6 (28.6%) | 15 (71.4%) | 7 (28.0%) | 18 (72.0%) |
| Grenada | 1 (16.7%) | 5 (83.3%) | 1 (25.0%) | 3 (75.0%) |
| Iran | 128 (12.5%) | 892 (87.5%) | 245 (10.5%) | 2082 (89.5%) |
| Iraq | 7 (14.3%) | 42 (85.7%) | 33 (15.8%) | 176 (84.2%) |
| Jamaica | 1 (6.7%) | 14 (93.3%) | 1 (11.1%) | 8 (88.9%) |
| Kazakhstan | 5 (17.2%) | 24 (82.8%) | 6 (14.6%) | 35 (85.4%) |
| Kyrgyzstan | 0 (0.0%) | 2 (100.0%) | 1 (50.0%) | 1 (50.0%) |
| Kuwait | 4 (5.5%) | 69 (94.5%) | 10 (14.5%) | 59 (85.5%) |
| Liberia | 1 (25.0%) | 3 (75.0%) | 1 (10.0%) | 9 (90.0%) |
| Montenegro | 0 (0.0%) | 5 (100.0%) | 1 (20.0%) | 4 (80.0%) |
| Mauritius | 1 (16.7%) | 5 (83.3%) | 1 (10.0%) | 9 (90.0%) |
| Malaysia | 37 (10.1%) | 331 (89.9%) | 69 (9.0%) | 700 (91.0%) |
| Nepal | 1 (12.5%) | 7 (87.5%) | 1 (3.8%) | 25 (96.2%) |
| Oman | 6 (12.0%) | 44 (88.0%) | 11 (12.1%) | 80 (87.9%) |
| Pakistan | 62 (27.8%) | 161 (72.2%) | 181 (21.5%) | 661 (78.5%) |
| Puerto Rico | 1 (2.9%) | 34 (97.1%) | 3 (12.5%) | 21 (87.5%) |
| Qatar | 16 (10.9%) | 131 (89.1%) | 23 (9.2%) | 228 (90.8%) |
| Saudi Arabia | 103 (15.3%) | 572 (84.7%) | 282 (16.7%) | 1402 (83.3%) |
| Sudan | 0 (0.0%) | 5 (100.0%) | 1 (14.3%) | 6 (85.7%) |
| Senegal | 4 (66.7%) | 2 (33.3%) | 5 (41.7%) | 7 (58.3%) |
| Trinidad &<br>Tobago | 0 (0.0%) | 13 (100.0%) | 1 (11.1%) | 8 (88.9%) |
| Tunisia | 6 (15.0%) | 34 (85.0%) | 8 (10.3%) | 70 (89.7%) |
| Vietnam | 9 (12.2%) | 65 (87.8%) | 20 (10.0%) | 181 (90.0%) |

## References

1. Hicks D, Wouters P, Waltman L, de Rijcke S, Rafols I. Bibliometrics: The Leiden Manifesto for research metrics. Nature. 2015 Apr;520(7548):429–31.

2. Ioannidis JPA, Baas J, Klavans R, Boyack KW. A standardized citation metrics author database annotated for scientific field. PLoS Biol. 2019;17(8):e3000384.

3. Ioannidis JPA, Klavans R, Boyack KW. Multiple Citation Indicators and Their Composite across Scientific Disciplines. PLoS Biol. 2016;14(7):e1002501.

4. Ioannidis JPA, Boyack KW, Baas J. Updated science-wide author databases of standardized citation indicators. PLoS Biol. 2020;18(10):e3000918.

5. Ioannidis JPA. Elsevier Data Repository, V6. 2023. October 2023 data-update for “Updated science-wide author databases of standardized citation indicators.”

6. Ioannidis JPA, Maniadis Z. In defense of quantitative metrics in researcher assessments. PLoS Biol. 2023;21(12):e3002408.

7. Ioannidis JPA, Maniadis Z. Quantitative research assessment: using metrics against gamed metrics. Intern Emerg Med. 2024 Jan;19(1):39–47.

8. Marcus A, Oransky I. Is There a Retraction Problem? And, If So, What Can We Do About It? In: Kathleen Hall Jamieson Dan M Kahan DAS, editor. The Oxford Handbook of the Science of Science Communication. Oxford University Press; 2017.

9. Oransky I. Retractions are increasing, but not enough. Nature. 2022;608(7921):9.

10. Oransky I. Volunteer watchdogs pushed a small country up the rankings. Science (1979). 2018;362(6413):395.

11. Hwang SY, Yon DK, Lee SW, Kim MS, Kim JY, Smith L, et al. Causes for Retraction in the Biomedical Literature: A Systematic Review of Studies of Retraction Notices. J Korean Med Sci. 2023;38(41).

12. Candal-Pedreira C, Ross JS, Ruano-Ravina A, Egilman DS, Fern’andez E, P’erez-R’os M. Retracted papers originating from paper mills: cross sectional study. BMJ. 2022;e071517.

13. Oransky I. Why misconduct could keep scientists from earning Highly Cited Researcher designations, and how our database plays a part [Internet]. 2022. Available from: https://retractionwatch.com/2022/11/15/why-misconduct-could-keep-scientists-from-earning-highly-cited-researcher-designations-and-how-our-database-plays-a-part/

14. Ioannidis JPA, Collins TA, Baas J. Evolving patterns of extreme publishing behavior across science. Scientometrics. 2024 Jul;1–4.

15. Hardwicke TE, Thibault RT, Kosie JE, Tzavella L, Bendixen T, Handcock SA, et al. Post-publication critique at top-ranked journals across scientific disciplines: A cross-sectional assessment of policies and practice. R Soc Open Sci. 2022 Aug 24;9(8).

16. Catanzaro M. Saudi universities entice top scientists to switch affiliations - sometimes with cash. Nature. 2023 May;617(7961):446–7.

17. Bhattacharjee Y. Citation impact. Saudi universities offer cash in exchange for academic prestige. Science. 2011 Dec;334(6061):1344–5.

18. Rodrigues F, Gupta P, Khan AP, Chatterjee T, Sandhu NK, Gupta L. The Cultural Context of Plagiarism and Research Misconduct in the Asian Region. J Korean Med Sci. 2023;38(12).

19. Rathore FA, Waqas A, Zia AM, Mavrinac M, Farooq F. Exploring the attitudes of medical faculty members and students in Pakistan towards plagiarism: a cross sectional survey. PeerJ. 2015 Jun;3:e1031.

20. Hsiao TK, Schneider J. Continued use of retracted papers: Temporal trends in citations and (lack of) awareness of retractions shown in citation contexts in biomedicine. Quantitative Science Studies. 2021;2(4):1144–69.

21. Marcus A, Abritis AJ, Oransky I. How to stop the unknowing citation of retracted papers. Anesthesiology. 2022;137(3):280–2.

22. Wager E, Williams P. Why and how do journals retract articles? An analysis of Medline retractions 1988-2008. J Med Ethics. 2011 Sep;37(9):567–70.

23. Baas J, Schotten M, Plume A, Côté G, Karimi R. Scopus as a curated, high-quality bibliometric data source for academic research in quantitative science studies. Quantitative Science Studies. 2020 Feb;1(1):377–86.

